# Sequence-to-function deep learning frameworks for synthetic biology

**DOI:** 10.1101/870055

**Authors:** Jacqueline Valeri, Katherine M. Collins, Bianca A. Lepe, Timothy K. Lu, Diogo M. Camacho

## Abstract

While synthetic biology has revolutionized our approaches to medicine, agriculture, and energy, the design of novel circuit components beyond nature-inspired templates can prove itself challenging without well-established design rules. Toehold switches — programmable nucleic acid sensors — face an analogous prediction and design bottleneck: our limited understanding of how sequence impacts functionality can require expensive, time-consuming screens for effective switches. Here, we introduce the Sequence-based Toehold Optimization and Redesign Model (STORM), a deep learning architecture that applies gradient ascent to re-engineer poorly-performing toeholds. Based on a dataset of 91,534 toehold switches, we examined convolutional filters and saliency maps of sequences to interpret our sequence-to-function model, identifying hot spots where mutations change toehold effectiveness and features unique to high-performing switches. Our modeling platform provides frameworks for future toehold selection, augmenting our ability to construct potent synthetic circuit components and precision diagnostics, and enabling straightforward translation of this *in silico* workflow to other circuitries.

## Introduction

Advances in synthetic biology have shifted paradigms in biotechnology by drawing inspiration from nature. While researchers have successfully isolated, integrated, and adapted templates from naturally occurring circuit parts — such as inducible promoters, terminators, and riboswitches — forward-engineering of novel components remains challenging. The workflow to develop a single circuitry may require weeks of screening and fine-tuning of individual units in efforts to perform an intended function as designed^1^. As such, there is a strong need for *in silico* screening of circuit parts in order to overcome prediction and design bottlenecks and ease integration of novel and redesigned synthetic components into existing biological systems.

In order to address the complexity of prediction and design of synthetic circuit parts, we focused on building computational tools to model nucleic acid sensors, such as riboswitches. Since the discovery of naturally occurring riboregulators — RNAs that alter their translation rate in the presence of small molecules and nucleic acids via changes in secondary structure^2,3^ — synthetic biologists have co-opted these circuit components for a variety of uses, from synthetic gene circuit construction^4,5^ to gene regulation^6,7^. The ability to engineer and control these molecules has broad applications spanning diverse areas, including medicine and agriculture^2,3^.

The toehold switch, a particularly versatile engineered riboregulator, is able to detect and respond to the presence of RNA molecules via linear-linear hybridization interactions^8^. A typical toehold switch anatomy consists of an unstructured toehold strand followed by a hairpin sequestering the Shine-Dalgarno sequence such that a reporter protein is not translated when the switch is in the OFF state (Fig. 1A, Table S1). The switch can be flexibly programmed such that the toehold and ascending stem of the hairpin are complementary to an arbitrary trigger RNA sequence, allowing the trigger RNA to hybridize to the toehold and subsequently melt the hairpin in the ON state, exposing the Shine-Dalgarno sequence to the ribosome and thereby initiating translation of a reporter gene. The inducible nature of toehold switches has led to their successful use as low-cost, freeze-dried, paper-based nucleic acid diagnostic tools^9–11^, as well as multiplexable components in complex genetic circuits with low crosstalk and high sensitivity^12–14^.

**Figure 1:**
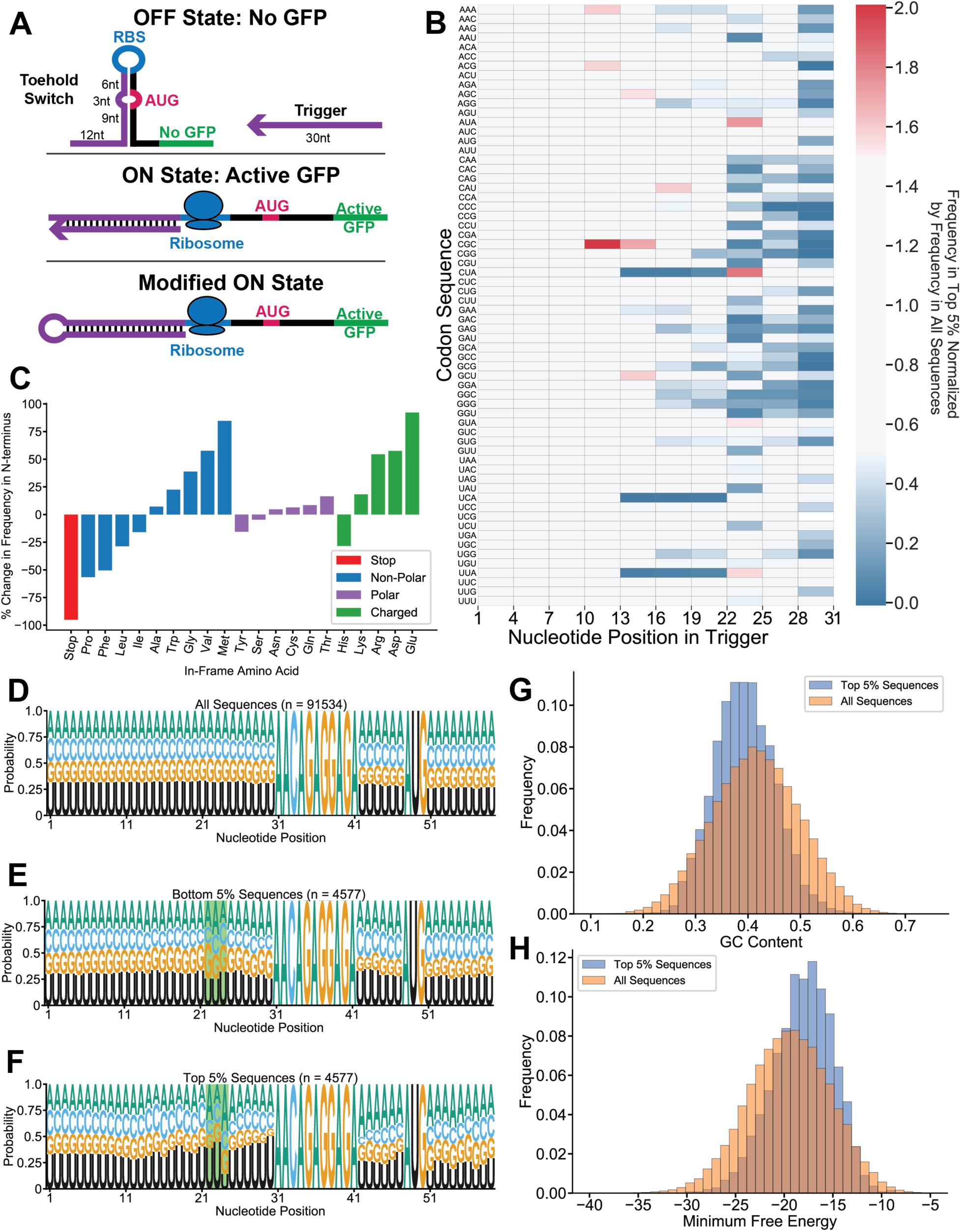
Top-performing toehold switches show over-representation of certain nucleotides at critical positions. (A) Toehold switches modify their secondary structure in response to the presence of a complementary RNA molecule known as a trigger. In absence of the trigger, the Shine-Dalgarno sequence, or ribosome binding site (RBS), remains inaccessible and the reporter protein is not translated (OFF state). Upon binding of the trigger to the switch, the hairpin melts, allowing ribosome recruitment to the Shine-Dalgarno sequence and subsequent translation of the downstream reporter GFP protein (ON state). A modified ON state switch was built for experimental testing so that one molecule could be tested with trigger and switch fused together. (B) Top-performing sequences were chosen by selecting the top 5% of toehold switches when ranked according to experimental ON/OFF ratios. Over-representation of each codon triplet at each position in the first 30 nucleotides of the switch was calculated by normalizing the frequency of the triplet in the top sequences by the frequency of the triplet across all sequences, with a normalized frequency higher than 1 indicating that the triplet is overrepresented in the top sequences. (C) Changes in frequencies of the in-frame amino acids located after the start codon in the N-terminus of the reporter protein were calculated by subtracting the frequency in the top 5% of sequences by the frequency in the bottom 5% and normalizing by the frequency in the bottom 5%. Sequence logos were calculated for the set of all sequences (D), the bottom 5% of sequences according to the experimental ON/OFF ratios (E), and the top 5% of sequences (F) by calculating the probability of each nucleotide occurring at each position in the switch sequence. GC content distributions (G) and switch minimum free energy estimates according to NUPACK^17^ (H) were calculated for all sequences and the top 5% of sequences, with clear differences in distributions (p < 5 × 10^-8^, Mann-Whitney U test).

Although toehold switches have become an effective and modular component of the synthetic biology toolkit, broad understanding of switch design has been limited by the small number of available toehold switches and lack of effective design rules for optimal performance. Sequence-based computational tools, which take into account thermodynamic equilibria and hybridization energies, have been developed to predict RNA sequence secondary structure prior to experimental validation^15–18^. However, when applied to multi-state toehold switches, these tools can lack predictive power and require time-intensive experimental screening, with correlations as low as 0.22 between predicted and measured efficacy^18^. An analysis of biophysical features conducted by Green et al.^8^ found only a few thermodynamic parameters that correlate to switch performance with an R^2^ value greater than 0.50, which are helpful heuristics but are not predictive of switch performance as standalone metrics. An improved prediction framework based on RNA sequence alone, customized for toehold switches and without reliance on biophysical parameters, would expand accessibility to efficiently tailor the design of these riboregulators for novel biological applications.

To improve toehold switch design and prediction, we took inspiration from the field of machine learning and deep learning. Machine learning approaches have been applied successfully to synthetic biology^1,19^, as exemplified in a recent study by Yang et al.^19^ using a ‘white box’ approach to extract antibiotic mechanisms of action, and in motif finding and DNA sequence prediction tasks^20–22^. As typical deep learning approaches require large amounts of training examples, we used a dataset from Angenent-Mari et al. (2019)^23^ that experimentally characterized 91,534 toehold switches, in both active and repressed states. We fed these switch sequences as input to a multi-layer convolutional neural network that can predict the ON and OFF state performances simultaneously. Our analysis provides both a deployable model to predict performance of new switches based on sequence alone, as well as a set of actionable design rules for optimal toehold switches. To understand the predictions made by the neural network, we used previously established white-box tools to examine motifs or partial motifs detected by the convolutional filters^24^, as well as positional importance of nucleotides. We provide examples of utilizing our deep neural network to rationally re-engineer poorly-performing toeholds using our Sequence-based Toehold Optimization and Redesign Model (STORM) framework, which employs gradient ascent to sample a sequence’s mutation space and customize toehold performance.

Sequence-to-function frameworks such as the one proposed here enable researchers to rapidly cycle through many possible design choices and select circuitries with optimal performance, while elucidating the underlying design principles for riboregulators. Given similarly sized datasets of other synthetic biology classes, translation of these frameworks to other training data and training tasks is both direct and desirable, and intersecting the cutting-edge features and techniques of both deep learning and synthetic biology holds profound implications for human health and biotechnology.

## Results

### Nucleotide over-representation in top-performing toehold sequences

A dataset of 244,000 toehold switches (Fig. 1A), including sequences tiled from viruses and the human genome as well as random sequences (see Methods), have been tested experimentally by Angenent-Mari et al^23^, with 91,534 switches meeting well-defined quality control criteria. Each toehold sequence is 59 nucleotides in length, with the first 30 nucleotides distinguishing the unique unstructured region followed by part of the hairpin; the remaining 29 nucleotides can be inferred by hairpin complementarity as well as Shine-Dalgarno sequence and start codon conservation (Table S1). Sequences with experimental ON/OFF ratios in the top and bottom 5^th^ percentile were characterized as high- and low-performing switches, respectively. To investigate nucleotide preference in high-performing switches, frequencies of triplets complementary to the trigger region were calculated with respect to the set of all sequences (Fig. 1B, Fig. S1). Of note is the over-representation of triplets AUA, CUA, GUA, and UUA at positions 22-24 in the switch, the three-nucleotide bulge directly opposite from the AUG start codon, suggesting that high-performing sequences may have an NUA at this bulge to prevent hybridization to the start codon (Fig. S2). Furthermore, the under-representation of UCA, CUA, and UUA at positions 13, 16, and 19 corresponds to the reverse complement of all three stop codons; positions 13 through 21 are opposite to the region directly following the start codon. Consistent with previous toehold design tools^18^, this indicates that the N-terminus of the reporter protein cannot tolerate an in-frame stop codon, which would terminate translation of the reporter protein.

To understand how changes to the coding part of the sequence (positions 51-59) affected toehold performance, we conducted a broader analysis of the in-frame amino acids at the N-terminus of the reporter gene (Fig. 1C, Fig. S3). In-frame stop codons were found less often at the N-terminus of high-performing sequences, in agreement with previous results. Additionally, though an unstructured linker region separates the switch from the reporter gene, we were nonetheless able to identify over- and under-representation of certain amino acids. Small hydrophobic amino acids such as valine, alanine, and glycine appear more often in the N-terminus of high-performing sequences than low-performing ones; top sequences also appear to contain less proline at the N-terminus, suggesting a slight preference for amino acids lacking a secondary amine; finally, polar amino acids appear less differentially represented than other classes, with differences in frequencies close to zero. Given these observations, toehold sequences appear sensitive to changes to in-frame amino acids at the N-terminus of the protein: small aliphatic amino acids are well-tolerated, but bulky non-polar amino acids such as phenylalanine and leucine should be avoided, if possible. However, due to the nature of the study, it is difficult to disentangle whether the observed enrichments for certain amino acids are due to structural changes at the RNA level or protein level, or perhaps due to differences in tRNA abundance.

Sequence logos based on positional probabilities of each nucleotide were calculated for all toeholds as well as for high-performing and low-performing toeholds separately to expand the nucleotide preference investigation to the entire 59-nucleotide switch sequence (Fig. 1D, E, F). Aside from the observed conservation of the Shine-Dalgarno (SD) and start codon sequences at positions 31-41 and 48-50, respectively, the sequence logo constructed from all 91,534 toeholds confirms that each of the four nucleotides are relatively evenly distributed at each position^23^. However, stratification into high- and low-performing sequences shows differential nucleotide composition immediately surrounding the SD sequence and following the start codon, with 51.75% of top-performing sequences containing uracil in the position immediately before the SD sequence and the same percentage of sequences containing adenine in the position immediately following the SD sequence. As we observed previously (Fig. 1B), the enrichment for NUA in positions 22-24 is highlighted in the top-performing logo with 16.1% of sequences containing the NUA motif: interestingly, 44.8% of sequences contain a U in position 22 and 44.7% of sequences contain an A in position 23, implying that single nucleotides belonging to the NUA motif are more prevalent than the complete motif. The sequence logo of poorly performing toehold suggests a corresponding NAU motif in positions 22-24, contributing evidence for a model that poorly functioning switches will hybridize successfully with the start codon, possibly preventing the hairpin from melting (Fig. S2).

### Top-performing switches are not adequately predicted by biophysical properties

The large size of the dataset allowed us to conduct an unbiased evaluation of toeholds’ biophysical properties suggested by other studies^15,18^. As previous reports have suggested that GC content is important for the strength of the ON and OFF state stabilities^8^, we compared the GC content distributions for top-performing sequences to that of all sequences (Fig. 1G), with a narrowing of the distribution and statistically significant shift to the left. These results suggest that successful toeholds may have a range of acceptable GC content between 20% and 60%. Very few top-performing sequences have more than 60% GC content, implying a necessity for A-U base pairing in the switch.

Multiple secondary structure prediction tools rely on thermodynamic modeling^15–18^; for example, the NUPACK software package calculates the equilibrium Gibbs free energy values for many possible secondary structures based on a provided RNA sequence, and reports to the user the most likely structure based on a minimum free energy (MFE) determination^15^. Because MFE has been quantified as an important predictive tool^8,15,17,18^, we investigated differences in predicted MFEs of toehold switches in our dataset by assessing the distribution in top-performing sequences. High-performing sequences had a statistically significantly higher MFE distribution than the set of all sequences (Fig. 1H), possibly due to extremely favorable MFE values indicating stability of the OFF state such that the hairpin will not melt in the presence of the trigger. As such, sequences with highly favorable free energy values should not be expected to perform better than sequences with less negative MFE values. Although top sequences exhibit statistically significant shifts in both GC content and MFE distributions, both properties lack predictive power due to their broad range of acceptable values.

### Multi-layer convolutional neural network to predict and classify toehold performance

As no single toehold property was found to be solely predictive of switch performance, a deep learning model was built to predict ON and OFF states based on switch sequence alone. Given the recent advancements in both accuracy and accessibility of deep learning^1,25^, a convolutional neural network (CNN)^26^ was constructed to take RNA sequences as input, employing two convolutional layers to identify motifs and partial motifs in the input sequences and their interactions^24^ (Fig. 2A). Following the convolutional layers, the model employs a multi-layer perceptron with three fully-connected layers, where every node in a given layer is connected to every node in the previous layer, to synthesize the features from the convolutions to output an ON and OFF prediction for each toehold sequence. As a comparison, neural networks with zero and one convolutional layers were built, and the performance of each model in a classification task — splitting sequences into the bottom 90% and top 10% of toehold switches — was compared via both a receiver operating characteristic (ROC) curve and a precision-recall (PR) curve in a 10-fold cross-validation setting (Fig. 2B, Fig. S4). The two-layer CNN ON classifier had an area-under-ROC curve (auROC) of 0.907, while the OFF classifier had an auROC of 0.868. To ensure that the 90% threshold was not arbitrarily effective for sequence classification, we tested an additional classifier built to stratify the top 50% and bottom 50% of sequences, which performed similarly (Fig. S5), with an auROC of 0.907 and 0.855 for the ON and OFF classifiers, respectively. The high auROC values suggest a high sensitivity for the classifier, while the high average precision of 0.979 to 0.989 for all models indicates that the number of false positives is low, both desirable traits for a classifier.

**Figure 2:**
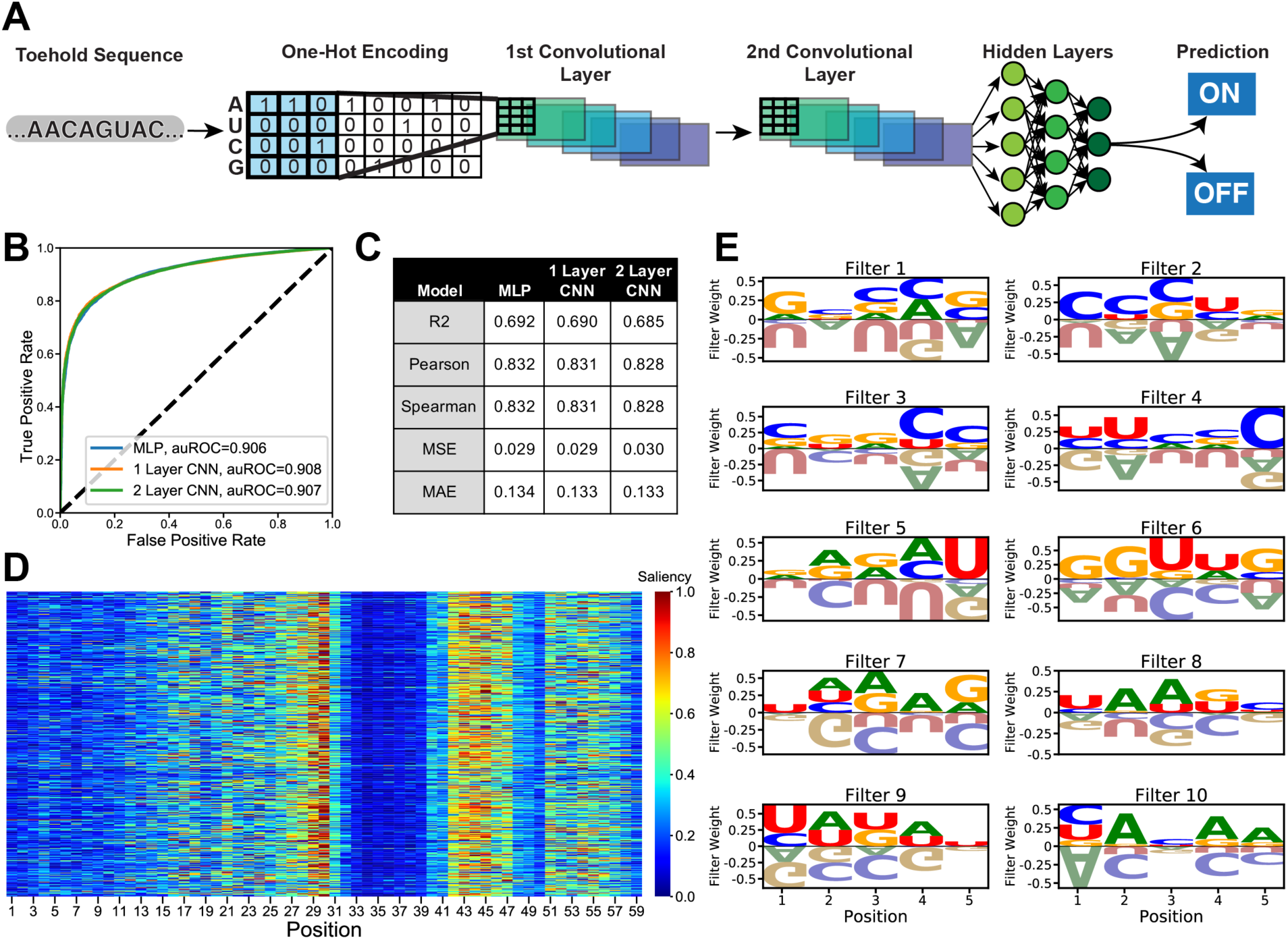
A convolutional neural network built to predict the ON and OFF values of toeholds switches can be examined with white-box tools. (A) A neural network was constructed that takes an RNA sequence as input and passes the one-hot encoded matrix to two convolutional layers, the first of which learns motifs or partial motifs in the sequences and the second of which may learn interactions between these motifs. The information is then passed to three fully connected layers and two output nodes, which can be used to predict ON and OFF values. The model, along with simplified models of one convolutional layer (1 Layer CNN) and zero convolutional layers (multi-layer perceptron, or MLP), were evaluated on a classification task: specifically, can we map toehold sequences into top performing (top 10^th^ percentile ON/OFF value) vs. poor performing (remaining 90^th^ percentile)? ROC and Precision-Recall curves were plotted for the 10-fold cross validation on both ON values (B, Fig. S4A) and OFF values (Fig. S4B, S4C). Regression model performance of ON (C) and OFF (Table S2) predictions were also compared between the MLP, 1 Layer CNN, and full-model 2 Layer CNN. To visualize how the model makes its decisions, 500 random sequences were selected and a saliency map (D) was plotted of the importance of each nucleotide to the model’s prediction of ON, with high saliency meaning the nucleotide at that position was critical to the model’s prediction. To understand the sequence patterns registered by the model, the learned filter weights for each of the 10 filters of width 5 in the first convolutional layer can be visualized as sequence logos similar to position weight matrices (E).

To expand the utility of the model, performance was also evaluated in a regression task to predict both ON (Fig. 2C) and OFF (Table S2) values directly. We evaluated all models on their R^2^ value, Pearson correlation coefficient, Spearman correlation coefficient (SCC), mean squared error (MSE), and mean absolute error (MAE). The best Pearson correlations achieved for ON and OFF, respectively, were 0.832 and 0.731, suggesting competent predictive ability of our model. Across all architectures, OFF predictions showed worse correlative metrics but better MAE and MSE metrics; for example, the two-layer CNN has an SCC of 0.828 for ON values, considerably outperforming the OFF SCC of 0.693. These results suggest that the models are able to learn features distinguishing a high ON value more easily than they can learn features distinguishing OFF states, possibly resulting from skewedness in the OFF data or from variance in the OFF state due to autofluorescence not being subtracted out^8^.

Although the two-layer CNN was not the top-performing model across all tasks, being surpassed slightly by a simpler multi-layer perceptron with zero convolutional layers, its multiple convolutional layers offer a level of interpretability beyond the other models showcased here. When CNNs are applied to images, consecutive convolutional layers learn additive features; for example, in facial analyses, edges and curves are learned in the first convolutional layer, followed by eyes, noses, and mouths, accumulating higher-order features through the layers^26^. As secondary structure is an important feature of toehold switches, we aimed to design a model such that the first convolutional layer learned motifs or partial motifs of the linear RNA sequence, and the second convolutional layer would learn additive or interactive features of such motifs, thereby improving the model accuracy through an implicit learning of secondary structure. Likewise, although the complete 59-nucleotide sequence contains some redundant information given complementarity in the hairpin, the 59-nucleotide sequence was used as input rather than the unique 30-nucleotide sequence so that the model could learn interactions within the linear switch sequence.

The regression model was used to predict the ON value of the 168 toehold switches reported in Green et al.^8^ in order to evaluate the model’s performance on unseen external data (Table S3). While the R^2^ is relatively low at 0.14, the higher Spearman correlation coefficient of 0.48 is an important characterization because a switch’s performance relative to another sequence can be more practically utilized than an arbitrary performance value. As an experimentalist may want to prioritize testing the best toehold out of a set of many, rank correlations may better inform decisions than precise ON/OFF ratios. Of note, no Green et al. switch sequences were included in our training data, suggesting that our models can be extended to unseen data without retraining the model or re-running the hyperparameter optimization, both computationally intensive and time-consuming steps.

### Improved interpretability of convolutional neural network predictions

Next, we harnessed a suite of white-box approaches to examine how our multi-layer CNN model was making its predictions. Given the vast number of weights and nonlinear functions that form the backbone of neural networks, it can be challenging to deduce why a model made the predictions it did^25^. While recent work, such as soft explainable decision trees^27^, have enabled researchers to look inside this ‘black-box’ by using a neural network to train a decision tree, we chose to visualize weights and activations of our trained model directly^28^. To understand the model’s predictions on an individual sequence, we took 500 random switches and evaluated the importance of each position in the sequence towards predicting the ON value (Fig. 2D); here, a higher saliency, computed by summing gradients across nucleotides at each position (see Methods), indicates that the nucleotide was considered to be more influential in the model’s ON prediction process. Our results show that swaths of unimportant regions mark the first twelve nucleotides in the sequence corresponding to the toehold region, as well as the constant ribosome binding site and the start codon, echoing results from codon and nucleotide representation in high-performing sequences (Fig. 1B, Fig. 2D). However, analogous to the sequence logos, the model interprets the area directly surrounding the Shine-Dalgarno sequence as being vital to toehold performance. Interestingly, position 30 appears more important than the nucleotide directly opposite it in the hairpin, position 42, indicating that the model does not need both positions to draw a conclusion about the ON value. To understand if the sequence saliency varies with the experimental values of the ON or OFF prediction, saliency maps for sets of high-, medium-, and poorly-performing toeholds were evaluated (Fig. S6). Poorly-performing toehold maps show similarly low activation in the first twelve nucleotides as their medium- and high-performing counterparts, suggesting that the model is not drawing conclusions by paying attention to different regions of toeholds. Conversely, these results may suggest that nucleotides in the ascending stem at positions 13-30 are important to characterize in all sequences regardless of performance.

Taking inspiration from work in the field of image recognition and genomics^26,29–31^, we investigated the first convolutional layer to see which features our model deemed important by interpreting the filter weights learned from input sequences as sequence logos (Fig. 2E). Exploration of the filter weights allowed us to identify a motif resembling the UAA stop codon in filters 7 and 8, though the filter does not explicitly correspond to any one region of the toehold and the direction of the effect of this motif on toehold performance is inconclusive. Filter 9 appears similar to the Pribnow box commonly found upstream of prokaryotic promoters, thereby indicating that significant information may be captured by some filters in our CNN. Further work is needed to explore the generalizability of these filters in the analyses of RNA sequences, and more definitive motif extraction could possibly be achieved by constructing an ensemble of convolutional networks and aggregating their convolutional filters for more statistical power.

### Rational redesign of toehold sequences by co-opting the pre-trained neural network

Given limitations in current toehold design processes, we pursued building a deep learning framework to rationally redesign sequences and optimize poorly-performing toeholds into high-performing ones via predictive point mutations. We converted our initial pre-trained model to build a Sequence-based Toehold Optimization and Redesign Model (STORM). Rather than using gradient descent as in the previous classification and regression tasks, we used gradient ascent to optimize sequences to meet target ON and OFF values (Fig. 3A, B). To evaluate the utility of STORM, we took the 100 worst experimental toeholds and engineered these to have a higher ON and lower OFF value, respectively (Fig. 3C, D). Clear differences emerged in the first ten to twelve nucleotides, with a run of cytosines that resemble the patterns learned by filters 2 and 4 of our two-layer CNN (Fig. 2E). The post-optimization sequences had an NUA motif enriched at positions 22-24, indicating the same preference for NUA that was seen in our previous triplet codon enrichment analysis (Fig. 1B, Fig. S2). These results confirm that the model can incorporate information learned in the training process to the redesign task, such that our framework is not limited to learning toehold sequence functionality, but rather, can be converted to other utilities such as toehold redesign. Saliency maps generated of the 100 worst sequences before and after optimization (Fig. S7) indicate that the model focuses more attention on the first ten to twelve nucleotides after optimization, signifying that the run of cytosines actionably changed the toehold performance prediction and were not an artifact of the redesign. Though the cytosine enrichment appears contradictory to results implying the first twelve nucleotides are inconsequential to model prediction, changes in model attention confirm that the cytosines contribute to model decision-making, encouraging the use of neural networks over simple nucleotide frequency studies which did not identify such a feature.

**Figure 3:**
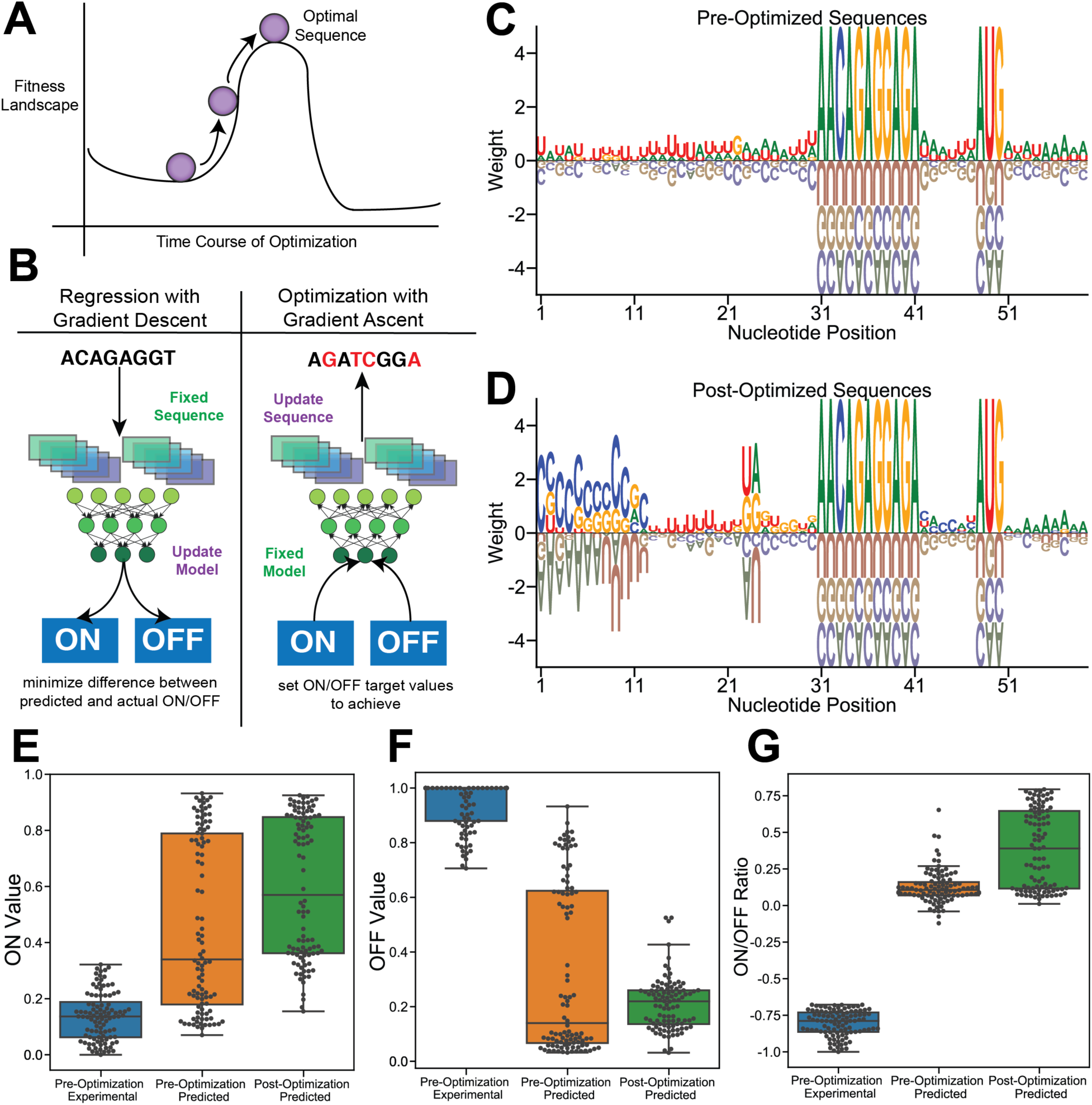
Poorly performing sequences can be transformed into highly performing toehold switches by co-opting the neural network for an optimization task. (A) Transformation of a poorly performing toehold switch can be achieved by ascending the fitness landscape of any given sequence to converge on a maxima in the landscape. (B) In contrast to the normal model optimization, which uses gradient descent to update the model and predict the ON and OFF values for any given sequence, the model can be inverted for gradient ascent of the sequence. Target ON and OFF values can be set, and a sequence is updated to achieve those target values while the trained model remains fixed. Position weight matrices can be plotted for a set of the 100 worst experimental toeholds both before (C) and after (D) optimization with gradient ascent; weight is the log_2_ of the probability of a nucleotide divided by the expected probability of the nucleotide. The ON values (E), OFF values (F), and ON/OFF ratios (G) can be plotted for the sequences before and after optimization.

For the set of 100 worst toeholds, the pre-optimization and post-optimization predicted ON and ON/OFF ratio values increase (Fig. 3E, G), which is expected given that the gradient ascent framework makes as many mutations to the sequence as possible to maximize the ON state while simultaneously minimizing the OFF state. Our results show that post-optimization predicted OFF values increase slightly from pre-optimization predicted OFF values (Fig. 3F), suggesting that for this set of optimized toeholds, higher performance was achieved by modifications to increase the ON value rather than decrease the OFF value. To ensure that these results were robust despite difficulties in predicting the ON and OFF values of experimentally poor toeholds, STORM was applied to a set of 100 toeholds with higher correlation in the experimental and predicted pre-optimization performance (Fig. S8). Interestingly, the focus on maximizing ON values was not observed in the set of 100 toehold sequences with a high correlation between their ON and OFF experimental and predicted values. These sequences had both high ON values and high OFF values at baseline, and the model focused on decreasing the OFF values rather than substantially increasing the ON values. However, the model’s OFF prediction quantified via R^2^ is less accurate than its ON prediction (Fig. 2C, Table S2), so it is unclear if this observation is an artifact arising from less reliable OFF prediction. Nevertheless, these results suggest that STORM is not following a static set of guidelines for optimization but rather adapts to each sequence by sampling from its custom mutation space. As the sequence optimization process may create a toehold that is not complementary with the original intended biological target, we envision STORM being utilized as a valuable tool for unconstrained sequence development, such as in synthetic circuit component construction.

## Discussion

Given the power of modular, programmable synthetic circuit components for diverse design problems and applications, there is a compelling need to better integrate computational and experimental approaches. We hoped to address this prediction and design bottleneck by building STORM, a deep learning framework that allows for the optimization of toehold switches according to user-inputted criteria. Our deployable models build on convolutional architectures and only require the RNA or DNA sequence of the trigger as input.

Our convolutional neural network offers meaningful insight into toehold design and pre-experimental performance prediction, underscoring the importance of designing models with interpretability in mind. Rather than treating the model as a black box and trusting its 15 predictions, recent advancements in machine learning have emphasized the importance of understanding how and why models reach their conclusions^28,32–34^. The convolutional neural network employed here gives us an opportunity to directly visualize the learned motifs of the network, highlighting potentially interesting biological features. Furthermore, saliency maps are a valuable tool to identify possible areas of model confusion and understand where the model focuses its attention^28,32^. Synthetic biology can thus benefit greatly from applying interpretable methods to deep learning frameworks.

Armed with a pre-trained convolutional neural network, our gradient ascent framework can be used to optimize any set of toehold sequences for any performance constraints. Though gradient ascent is not a new concept^35^, the application of generative models to redesign linear sequences for the end goal of improving function has been gaining traction in protein engineering^36,37^. For instance, generative adversarial networks^38^ (GANs) — a modeling paradigm that simultaneously trains two competing neural networks — are being used to teach a network to produce realistic protein structure maps^39^, in the same way that such tools have been used in computer vision to make realistic celebrity faces^40^. However, GANs remain challenging to train and define for biological tasks. By comparison, STORM readily converts our existing predictive CNN without extensive re-training, allowing for the redesign of sub-optimal toehold switches in a generative manner. Given more popular and accurate sequence-to-function prediction models^41^, STORM could be used as a guide for new nucleic acid modeling problems such as combinatorial circuit design, as well as to look inside of and augment existing prediction frameworks.

Additionally, the STORM framework allowed us to concertedly identify key design rules for this class of riboregulators. Importantly, both high-performing toeholds and optimized redesigned sequences have an over-representation of NUA at positions 22-24, suggesting that toehold performance suffers if there is hybridization with the start codon. Additionally, bulky hydrophobic amino acids or amino acids that disrupt polypeptide structures are unfavorable at the N-terminus, with charged amino acids appearing over-represented in top sequences. The first twelve nucleotides in the sequence corresponding to the toehold region exhibit evidence of being less important than the remaining nucleotides, indicating that the first twelve nucleotides can be freely designed based on trigger complementarity without constraining any design rules. However, the STORM redesign preferentially transformed the first twelve nucleotides to cytosines, implying that particularly well-designed arrangements of nucleotides in these positions can affect toehold performance. While GC content and MFE of the sequence should be considered, these properties are not strongly predictive of experimental performance, but can be included as additional considerations in switch selection.

It is important to note that the tools developed here are not constrained to any single riboregulator design or dataset. Our neural network architecture can be adapted for any RNA or DNA sequence with a measurable performance, dependent only on a large enough set of data to perform model training. Similarly, white-box tools are model-agnostic, with applicability to any convolutional model and nucleic acid dataset. Our gradient ascent framework to transform poorly performing toehold sequences can be applied to any set of toeholds or similar nucleic acid sequences. With the advent of tools to design, test, and process high-throughput biological datasets, machine learning could be fully exploited as a means to glean new insight into synthetic biology components, tools, and phenomena.

## Methods

### Toehold Sequence Generation

In order to define the sequences to be tested (see Angenent-Mari et al.^23^), we perform a sequence tiling on a variety of prokaryotic genomes, as well as the complete set of human transcriptional regulators. Briefly, each chosen sequence was tiled with a sequence length of 30 nucleotides and a stride of 5 nucleotides. Additionally, we generated a set of 10,000 random sequences of length 30, drawing each nucleotide from a uniform distribution at each position. This approach resulted in a set of 244,000 sequences to be synthesized and tested experimentally.

### Data Filtering and Visualization

244,000 toehold sequences were tested by Angenent-Mari et al.^23^ and the experimental data was obtained as logarithm-transformed GFP fluorescence measured at both the modified ON state (with trigger present, fused to the switch sequence) and OFF state (without trigger). Measurements were normalized and quality control was performed as indicated in Angenent-Mari et al^23^, resulting in 91,534 sequences. Additionally, a final filtering step was applied prior to training the neural network, where sequences were split into 1000 bins for both ON and OFF distributions, and bins were down-sampled to the mean number of counts across all bins (Fig. S9). The union of sequences from both the ON and OFF filtering stage was carried forward, resulting in 81,155 switches. All sequence logos were visualized with LogoMaker^42^.

### Model Architecture

The model was constructed of two convolutional layers to detect genomic motifs^24,28,43,44^. The first convolutional layer consisted of 10 filters of width 5; the second convolutional layer consisted of 5 filters of width 3. The filters, or weight matrices, were convolved over nucleotide channels and point-wise multiplied with the input sequence, with the magnitude of this multiplication, or activation, corresponding to the degree of similarity between the filter pattern and the input^28^. Activations from the second convolutional layer were flattened into a one-dimensional vector and fed as input to three fully connected layers with successively decreasing numbers of nodes (150, 60, 15, respectively). All layers applied the rectified linear unit (ReLU) nonlinearity function to node outputs^25^ and these activations were passed independently to two output layers: the ON and OFF prediction outputs, respectively. For the regression task, the last fully connected layer utilized linear activation to output continuous ON and OFF values; in contrast, a softmax activation was used for the classification task in order to ensure normalized class probabilities. After each layer, a dropout rate^45^ of 0.3 was applied, and the ridge regression (L2 regularization) coefficient on the activations was set to 0.001 to increase the sparsity of the hidden layers and decrease the risk of overfitting by constraining the values of the weights^25^. Errors between true and predicted ON and OFF values were computed over small batches of 128 toehold sequences at a time using a binary cross-entropy loss function. This loss information was backpropagated through the model, and stochastic gradient descent via the Adam optimizer was used to update the weights of the model such that the disparity between model predictions and true value was minimized^25^. Adam was used with a learning rate of 0.005 to train the model at a speed that achieved fast convergence without over- or under-shooting the optimal model fit. Weights were updated with respect to both ON prediction and OFF prediction simultaneously. Keras with a TensorFlow^46^ backend was used to construct and optimize the model.

### Hyperparameter Optimization via Grid Search

A grid search was conducted to find the best model structure and architecture settings (Supplemental Information File 1). We explored within the space of convolutional architectures, as recurrent neural networks (RNNs) tend to be harder to disentangle why predictions were made, and CNN structures have grown increasingly popular for sequence-based tasks^43^. Architecture design parameters were selected randomly rather than combinatorically as in a traditional grid search to enable a broader search of the architecture landscape within time and computation constraints^47^. The convolutional hyperparameters were varied to maximize the convolutional layers’ ability to learn a sufficient number of short, meaningful patterns of relevant nucleotide combinations^24,44^. In accordance with Occam’s Razor, model simplicity was prioritized to avoid unnecessary complexity (i.e. more convolutional filters than needed) that did not augment the biological interpretability of the model^48^.

Five-fold cross validation was used to train and evaluate each parameter combination. For each fold, and for ON and OFF predictions separately, R^2^ and Spearman’s rank correlation were calculated to estimate the generalizability of the model. The best architecture was selected by sorting the results by their combined average R^2^ over ON and OFF prediction and choosing an architecture with first layer filters that would enable downstream interpretation of biologically-meaningful motifs and maximal ability to decode predictions in the context of toehold design rules.

### Model Evaluation

To assess the best architecture performance and train the final model, the data was shuffled and iteratively split using 10-fold cross validation; the test set per fold was further split in half to be used as validation toehold sequences to select the optimal number of training epochs. A stratified split method enabled the cross-validation to be conducted with the class imbalance from the 90%/10% classification split preserved in each fold. A deployable model was trained on 85% of the data, with 15% held-out as validation data to enable early stopping in training. The same architecture found in the grid search was used for the binary classification task and trained using the 10-fold cross validation procedure detailed above; the area-under-ROC curve values and area-under-PR curve values were calculated for ON and OFF classification tasks.

### Saliency Maps

Saliency maps were generated to visualize which positions and nucleotides mattered most towards high ON and low OFF model predictions^28,32,33^. The keras-vis package was used to analyze how small changes in a given input toehold sequence change the model’s output predictions. The gradients were computed to highlight changes to the input sequence that produce large changes in the output predictions, revealing which positions in the toehold sequence were prioritized the most when predicting ON and OFF values. A saliency (i.e., an “importance score”) for each position and each nucleotide at that position was calculated by summing the gradients across all positions and nucleotides for each toehold. This saliency was normalized by the number of times a given nucleotide appeared at that position to control for more frequent nucleotides.

### STORM: A Sequence-based Toehold Optimization and Redesign Model

We converted our predictive pipeline to redesign poorly performing toehold switches via gradient ascent. The 100 toeholds with the lowest ON/OFF ratio were one-hot encoded and fed as inputs to the static model. Target ON and OFF values of 0.99 and 0.001, respectively, were set and supplied to SeqProp, an open-source python package that enables streamlined development of gradient ascent pipelines for genomic and RNA biology applications^22^. Toehold design constraints were incorporated into the loss function, such that the modified toehold switch contained the conserved sequences and base pairing within the hairpin was preserved. At each iteration, the ON and OFF values of the initial toehold sequence were predicted and the difference between the predicted values and target values was computed. This discrepancy between predicted and target values was then propagated back through the model to update the input sequence in the direction that decreased the difference between the predicted ON and OFF values and the target. The updated toehold position weight matrix was used as input to the next round of optimization, and at the last round of iteration, the final sequence was composed of nucleotides with the highest probabilities in the position weight matrix.

## Acknowledgements

We thank Nico Angenent-Mari and Luis R. Soenksen for feedback on model construction and for sharing with us the experimental data on the toehold designs, and Miguel A. Alcantar for help conceptualizing the gradient ascent framework. We also thank Amy Xiao, Megan Tse, and Pablo Cárdenas for reviewing figures and text. We thank Bogard et al. for help adapting their SeqProp framework for toehold switches. We thank James J. Collins for insightful critique of our work.

## Authors contributions

J.V. and K.M.C. designed the neural network architecture, visualized the model predictions, and engineered the gradient ascent framework. B.A.L. analyzed random grid search results and evaluated Green et al. sequences. T.K.L. and D.M.C. directed research and provided manuscript feedback. All authors wrote the manuscript.

## Data and materials availability

Request for data and code should be addressed to D.M.C. The Python code for neural net construction, training, and prediction can be found at this GitHub repository: www.github.com/midas-wyss/storm

## Supplementary Materials

Methods

Figures S1-S9

Tables S1-S3

Supplementary File 1

References (42-48)

## References

1. Camacho, D. M., Collins, K. M., Powers, R. K., Costello, J. C. & Collins, J. J. Next-Generation Machine Learning for Biological Networks. Cell 173, 1581–1592 (2018).

2. Hallberg, Z. F., Su, Y., Kitto, R. Z. & Hammond, M. C. Engineering and In Vivo Applications of Riboswitches. Annu. Rev. Biochem. 86, 515–539 (2017).

3. Serganov, A. & Nudler, E. A Decade of Riboswitches. Cell 152, 17–24 (2013).

4. Callura, J. M., Dwyer, D. J., Isaacs, F. J., Cantor, C. R. & Collins, J. J. Tracking, tuning, and terminating microbial physiology using synthetic riboregulators. Proc. Natl. Acad. Sci. 107, 15898–15903 (2010).

5. Rodrigo, G., Landrain, T. E. & Jaramillo, A. De novo automated design of small RNA circuits for engineering synthetic riboregulation in living cells. Proc. Natl. Acad. Sci. 109, 15271–15276 (2012).

6. Isaacs, F. J. et al. Engineered riboregulators enable post-transcriptional control of gene expression. Nat. Biotechnol. 22, 841–847 (2004).

7. Mutalik, V. K., Qi, L., Guimaraes, J. C., Lucks, J. B. & Arkin, A. P. Rationally designed families of orthogonal RNA regulators of translation. Nat. Chem. Biol. 8, 447–454 (2012).

8. Green, A. A., Silver, P. A., Collins, J. J. & Yin, P. Toehold Switches: De-Novo-Designed Regulators of Gene Expression. Cell 159, 925–939 (2014).

9. Pardee, K. et al. Paper-Based Synthetic Gene Networks. Cell 159, 940–954 (2014).

10. Pardee, K. et al. Rapid, Low-Cost Detection of Zika Virus Using Programmable Biomolecular Components. Cell 165, 1255–1266 (2016).

11. Ma, D., Shen, L., Wu, K., Diehnelt, C. W. & Green, A. A. Low-cost detection of norovirus using paper-based cell-free systems and synbody-based viral enrichment. Synth. Biol. 3, (2018).

12. Takahashi, M. K. & Lucks, J. B. A modular strategy for engineering orthogonal chimeric RNA transcription regulators. Nucleic Acids Res. 41, 7577–7588 (2013).

13. Green, A. A. et al. Complex cellular logic computation using ribocomputing devices. Nature 548, 117–121 (2017).

14. Kim, J. et al. De-Novo-Designed Translational Repressors for Multi-Input Cellular Logic. bioRxiv 501783 (2018) doi: 10.1101/501783.

15. Zadeh, J. N., Wolfe, B. R. & Pierce, N. A. Nucleic acid sequence design via efficient ensemble defect optimization. J. Comput. Chem. 32, 439–452 (2011).

16. Lorenz, R. et al. ViennaRNA Package 2.0. Algorithms Mol. Biol. AMB 6, 26 (2011).

17. Salis, H. M. Chapter two - The Ribosome Binding Site Calculator. in Methods in Enzymology (ed. Voigt, C.) vol. 498 19–42 (Academic Press, 2011).

18. To, A. C.-Y. et al. A comprehensive web tool for toehold switch design. Bioinformatics 34, 2862–2864 (2018).

19. Yang, J. H. et al. A White-Box Machine Learning Approach for Revealing Antibiotic Mechanisms of Action. Cell 177, 1649-1661.e9 (2019).

20. Singh, S. & Singh, R. Application of supervised machine learning algorithms for the classification of regulatory RNA riboswitches. Brief. Funct. Genomics 16, 99–105 (2017).

21. Hiscock, T. W. Adapting machine-learning algorithms to design gene circuits. BMC Bioinformatics 20, 214 (2019).

22. Bogard, N., Linder, J., Rosenberg, A. B. & Seelig, G. A Deep Neural Network for Predicting and Engineering Alternative Polyadenylation. Cell 178, 91-106.e23 (2019).

23. Angenent-Mari, N., Garruss, A. & Soenksen, L. Deep learning for RNA synthetic biology. Submitted for publication (2019).

24. Koo, P. K. & Eddy, S. R. Representation Learning of Genomic Sequence Motifs with Convolutional Neural Networks. bioRxiv 362756 (2019) doi: 10.1101/362756.

25. LeCun, Y., Bengio, Y. & Hinton, G. Deep learning. Nature 521, 436–444 (2015).

26. LeCun, Y., Bottou, L., Bengio, Y. & Ha, P. Gradient-Based Learning Applied to Document Recognition. 46 (1998).

27. Frosst, N. & Hinton, G. Distilling a Neural Network Into a Soft Decision Tree. 171109784 Cs Stat (2017).

28. Zeiler, M. D. & Fergus, R. Visualizing and Understanding Convolutional Networks. 13112901 Cs (2013).

29. Cuperus, J. T. et al. Deep learning of the regulatory grammar of yeast 5′ untranslated regions from 500,000 random sequences. Genome Res. 27, 2015–2024 (2017).

30. Movva, R. et al. Deciphering regulatory DNA sequences and noncoding genetic variants using neural network models of massively parallel reporter assays. PLOS ONE 14, e0218073 (2019).

31. Zou, J. et al. A primer on deep learning in genomics. Nat. Genet. 51, 12–18 (2019).

32. Simonyan, K., Vedaldi, A. & Zisserman, A. Deep Inside Convolutional Networks: Visualising Image Classification Models and Saliency Maps. 13126034 Cs (2013).

33. Shrikumar, A., Greenside, P. & Kundaje, A. Learning Important Features Through Propagating Activation Differences. 170402685 Cs (2017).

34. Budach, S. & Marsico, A. pysster: Classification of Biological Sequences by Learning Sequence and Structure Motifs with Convolutional Neural Networks. http://biorxiv.org/lookup/doi/10.1101/230086 (2017) doi: 10.1101/230086.

35. Erhan, D., Bengio, Y., Courville, A. C. & Vincent, P. Visualizing Higher-Layer Features of a Deep Network. in (2009).

36. Yu, M. K. et al. Visible Machine Learning for Biomedicine. Cell 173, 1562–1565 (2018).

37. Yang, K. K., Wu, Z. & Arnold, F. H. Machine-learning-guided directed evolution for protein engineering. Nat. Methods 16, 687–694 (2019).

38. Goodfellow, I. et al. Generative Adversarial Nets. in Advances in Neural Information Processing Systems 27 (eds. Ghahramani, Z., Welling, M., Cortes, C., Lawrence, N. D. & Weinberger, K. Q.) 2672–2680 (Curran Associates, Inc., 2014).

39. Anand, N. & Huang, P. Generative modeling for protein structures. in Advances in Neural Information Processing Systems 31 (eds. Bengio, S. et al.) 7494–7505 (Curran Associates, Inc., 2018).

40. Karras, T., Aila, T., Laine, S. & Lehtinen, J. PROGRESSIVE GROWING OF GANS FOR IMPROVED QUALITY, STABILITY, AND VARIATION. 26 (2018).

41. AlQuraishi, M. End-to-End Differentiable Learning of Protein Structure. Cell Syst. 8, 292-301.e3 (2019).

42. Tareen, A. & Kinney, J. B. Logomaker: Beautiful sequence logos in python. bioRxiv 635029 (2019) doi: 10.1101/635029.

43. Lai, G., Chang, W.-C., Yang, Y. & Liu, H. Modeling Long- and Short-Term Temporal Patterns with Deep Neural Networks. in The 41st International ACM SIGIR Conference on Research & Development in Information Retrieval - SIGIR ‘18 95–104 (ACM Press, 2018). doi: 10.1145/3209978.3210006.

44. Zeng, H., Edwards, M. D., Liu, G. & Gifford, D. K. Convolutional neural network architectures for predicting DNA–protein binding. Bioinformatics 32, i121–i127 (2016).

45. Srivastava, N., Hinton, G., Krizhevsky, A., Sutskever, I. & Salakhutdinov, R. Dropout: A Simple Way to Prevent Neural Networks from Overfitting. 30.

46. Abadi, M. et al. TensorFlow: Large-Scale Machine Learning on Heterogeneous Distributed Systems. 160304467 Cs (2016).

47. Bergstra, J. & Bengio, Y. Random Search for Hyper-Parameter Optimization. 25.

48. Domingos, P. The Role of Occam’s Razor in Knowledge Discovery. 19.

