## Supplement for "Sequence-to-function deep learning frameworks for synthetic biology"

**Supplemental File 1: Model performance was optimized via a hyperparameter random grid search.** To find the optimal model architecture, several hyperparameters were varied and randomly sampled. The number of filters tested in the first layer were 5, 10, and 15, testing each combination with filter widths of 3, 5, or 7. The number of filters tested in the second layer were 5 and 10, with filter widths of 3, 5, or 7. Additional hyperparameters such as dropout rate, L2 regularization, and Adam optimizer learning rate were varied to curtail overfitting and modify how the space of weights was explored during training: the dropout rates tested were 0.1, 0.3, and 0.5; L2 regularization values tested were 0 and 0.0001; and learning rate values tested were 0.0005 and 0.001. For all hyperparameter combinations, $R^2$ and Spearman correlation coefficients were evaluated across both ON and OFF values to ensure model predictions are sufficiently consistent with experimental results.

### Figures and Tables

| Component | Start Position | End Position | Length (nts) | Complementary to |
|---|---|---|---|---|
| Unstructured Region | 1 | 12 | 12 | Trigger (19-30) |
| Ascending Stem 1 | 13 | 21 | 9 | Trigger (10-18), Descending Stem 2 |
| Bulge Opposite AUG | 22 | 24 | 3 | |
| Ascending Stem 2 | 25 | 30 | 6 | Trigger (1-6), Descending Stem 1 |
| Shine-Dalgarno / Ribosome Binding Site (RBS) | 31 | 41 | 11 | |
| Descending Stem 1 | 42 | 47 | 6 | Ascending Stem 2 |
| AUG | 48 | 50 | 3 | |
| Descending Stem 2 | 51 | 59 | 9 | Ascending Stem 1 |

**Supplemental Table 1: Toehold switch anatomy consists of an unstructured region followed by a hairpin.** The hairpin stem relies on complementary nucleotides to maintain secondary structure. The loop of the hairpin contains the Shine-Dalgarno sequence, which becomes accessible to a ribosome when the 30-nucleotide trigger binds to the complementary switch region and melts the hairpin.

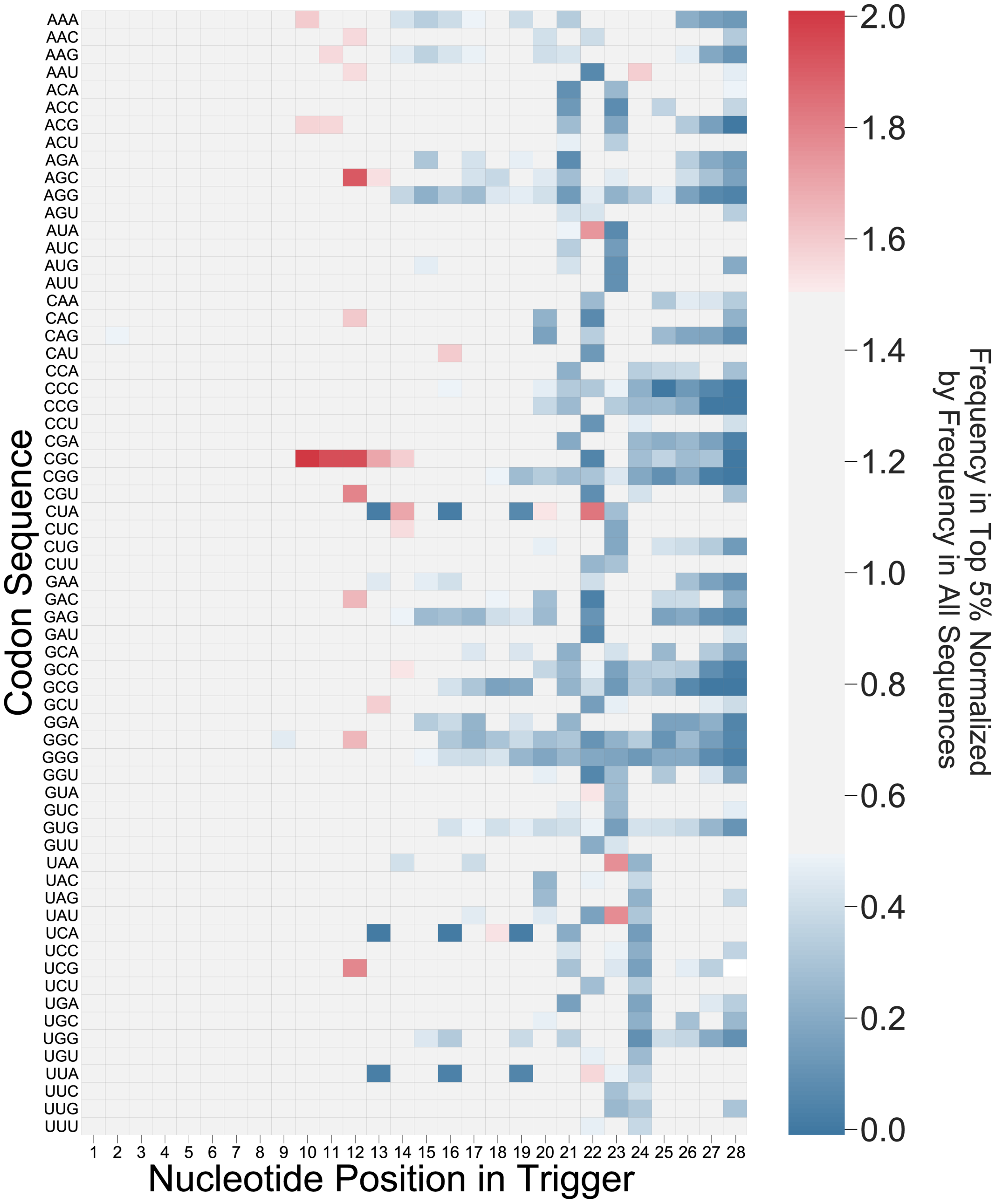


**Supplemental Figure 1: Top-performing sequences display preferences for certain codons at crucial positions.** The ratio of every codon triplet at each position in the trigger was calculated by dividing the frequency in the top 5% of sequences by the frequency in all sequences. The over-representation of NUA at position 22 is still visible, as in Fig. 1B. The under-representation of CUA, UCA, UUA — reverse complements of the start codons — is restricted to in-frame positions 13, 16, and 19, which correspond to the N-terminus of the translated reporter protein.

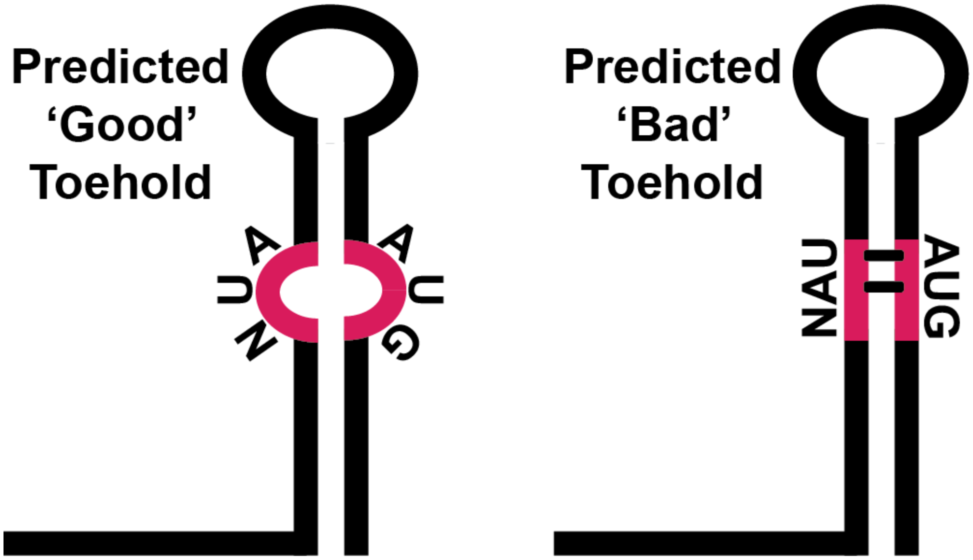


**Supplementary Figure 2: NUA is over-represented in the 3-nucleotide bulge opposite the start codon.** High-performing toeholds have an over-representation of NUA in positions 22-24, the 3-nucleotide bulge opposite the start codon, suggesting that top-performing sequences do not hybridize with the start codon.

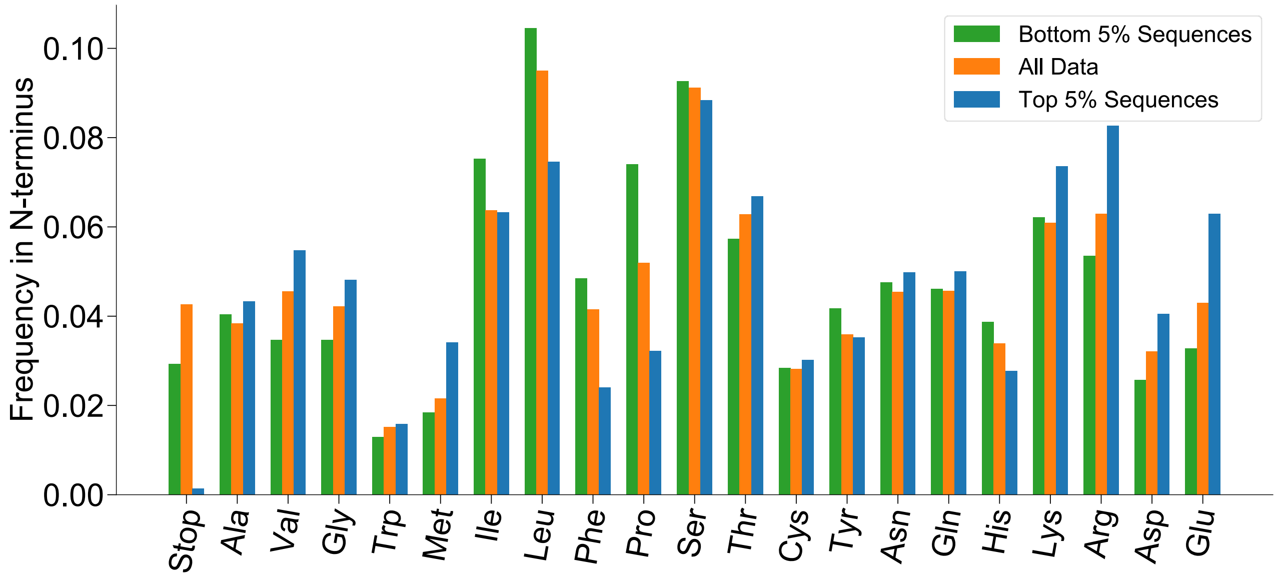


**Supplementary Figure 3: High-performing and poorly-performing switches show over-representation for specific types of amino acids in the N-terminus.** Raw frequencies of each in-frame amino acid were calculated for all sequences, as well as for the top and bottom 5% of sequences when stratified according to experimental ON/OFF ratios. As in Fig. 1C, under-representation of stop codons, phenylalanine, and proline is evident in top sequences, while top sequences seem to prefer charged amino acids in the N-terminus.

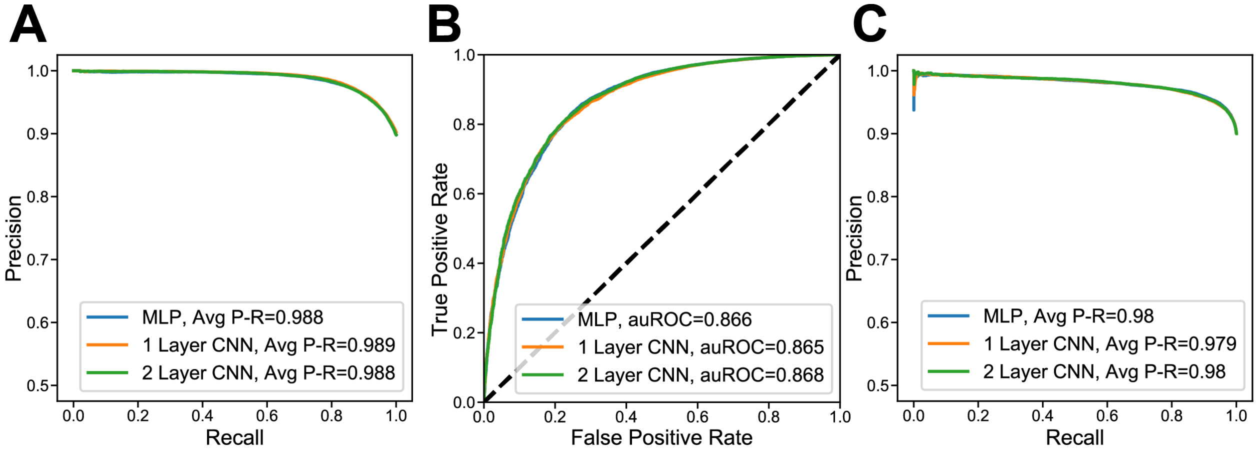


**Supplementary Figure 4: The convolutional neural network classifies toeholds with a low rate of fall positives and false negatives for both ON and OFF values.** The precision-recall (P-R) curve (Fig. S4A) shows both high precision and recall for the classification of the top 10%/bottom 90% of ON values, complementing the ROC curve in Fig. 2B. The classification of OFF values into top 10%/bottom 90% performs similarly, with a slight reduction in accuracy and precision: the ROC curve (Fig. S4B) and P-R curve (Fig. S4C) for OFF values both have high area under the curve values.

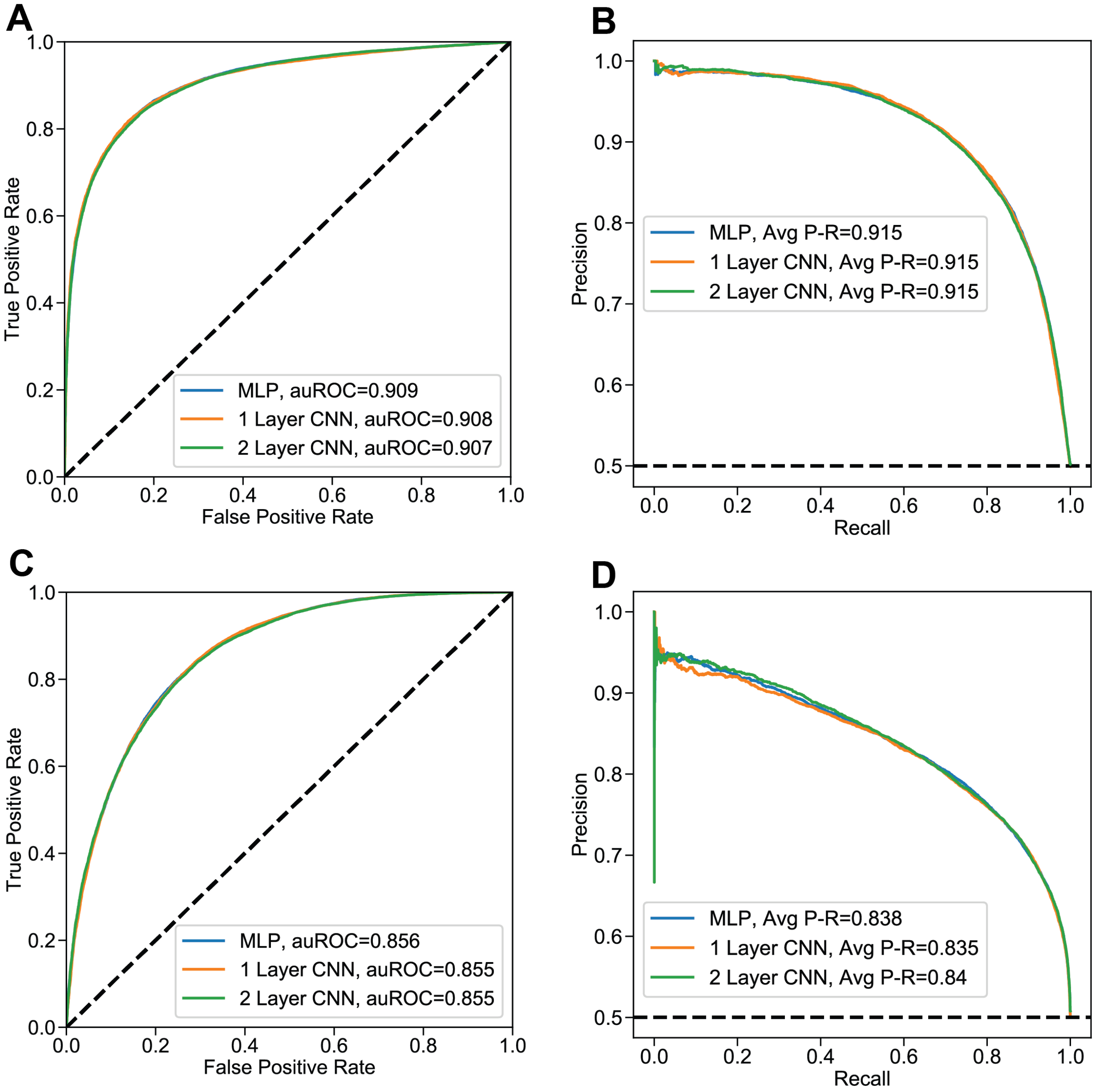


**Supplemental Figure 5: Thresholding on median ON and OFF values for classification performs similarly to 90$^{th}$ percentile threshold classification.** Rather than use the 90$^{th}$ percentile as a threshold for toehold classification as in Fig. 2B and Fig. S4, the median ON and OFF values respectively were used to ensure that the model was robust enough to accurately stratify the top half and bottom half of sequences even when changing the classification threshold. Both ON (Fig. S5A, B) and OFF (Fig. S5C, D) classification accuracy was measured with receiver operating characteristic (ROC) and precision-recall (P-R) curves, with only a slight reduction in the area under the ROC curve (auROC) and average precision-recall from the 90$^{th}$ percentile classification.

| Model | MLP | 1 Layer CNN | 2 Layer CNN |
|---|---|---|---|
| R2 | 0.529 | 0.535 | 0.524 |
| Pearson | 0.727 | 0.731 | 0.724 |
| Spearman | 0.696 | 0.693 | 0.693 |
| MSE | 0.024 | 0.024 | 0.024 |
| MAE | 0.113 | 0.113 | 0.114 |

**Supplemental Table 2: The convolutional neural network performs well when predicting continuous values for the OFF state.** The regression of OFF values performs similarly to ON value regression (Fig. 2C). Analogous to the classification task, most OFF metrics are slightly worse than the ON metrics, with the except of mean squared error (MSE) and mean absolute error (MAE), suggesting that predictions of ON and OFF states may have differing performances based on intrinsic properties of the ON and OFF distributions.

| | $R^2$ | Spearman | Pearson |
|---|---|---|---|
| Statistic | 0.1408 | 0.4782 | 0.3752 |
| P-Value | 5.42E-07 | 5.53E-11 | 5.42E-07 |

**Supplemental Table 3: Green et al. toehold switches validate the model without re-training or re-optimizing model architecture**. To assess the correlation between the normalized ON values from the set of 168 toehold switches tested in Green et. al. and the predicted ON values from our regression model, we calculated the $R^2$ value, Spearman correlation coefficient, and Pearson correlation coefficient. Of particular interest, the Spearman correlation coefficient is significantly higher than what would be expected if there was no relationship between the normalized experimental ON and the predicted ON values (i.e., Spearman correlation of zero).

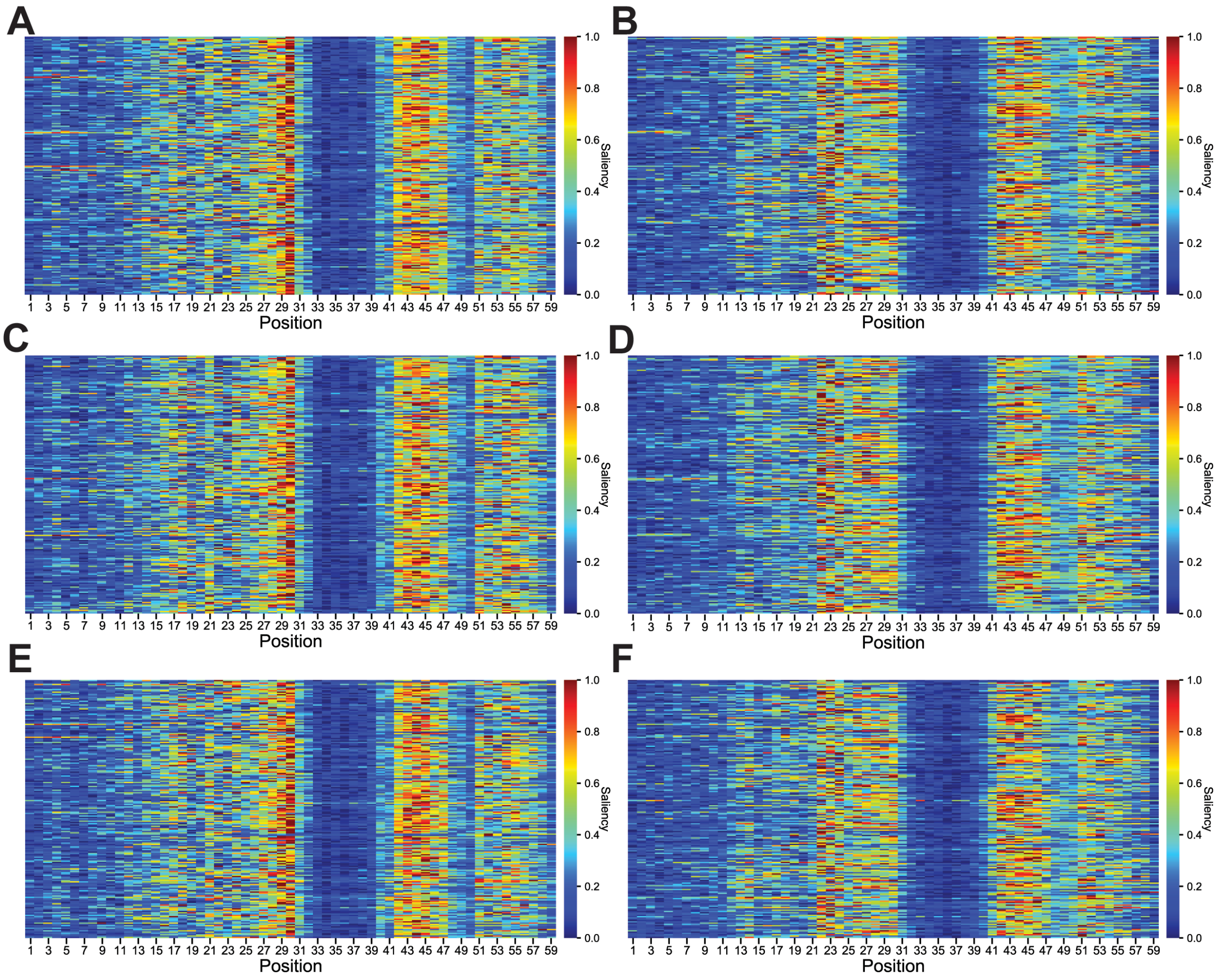


**Supplemental Figure 6: Saliency maps are not markedly different for toehold switches with high, medium, and poor performance.** To ensure that the saliency maps did not change dramatically based on which sequences were tested, saliency maps were generated for sets of ‘good’ (Fig. S6A, B), ‘medium’ (Fig. S6C, D), and ‘bad’ (Fig. S6E, F) switches according to ON/OFF ratios. For each set of sequences, both a saliency map to illustrate the parts of the sequence that maximized the ON prediction (Fig. S6A, C, E) as well as a saliency map to illustrate the parts of the sequence that minimized the OFF prediction (Fig. S6B, D, F) were generated. All saliency maps indicate the importance of the regions immediately surrounding the Shine-Dalgarno sequence.

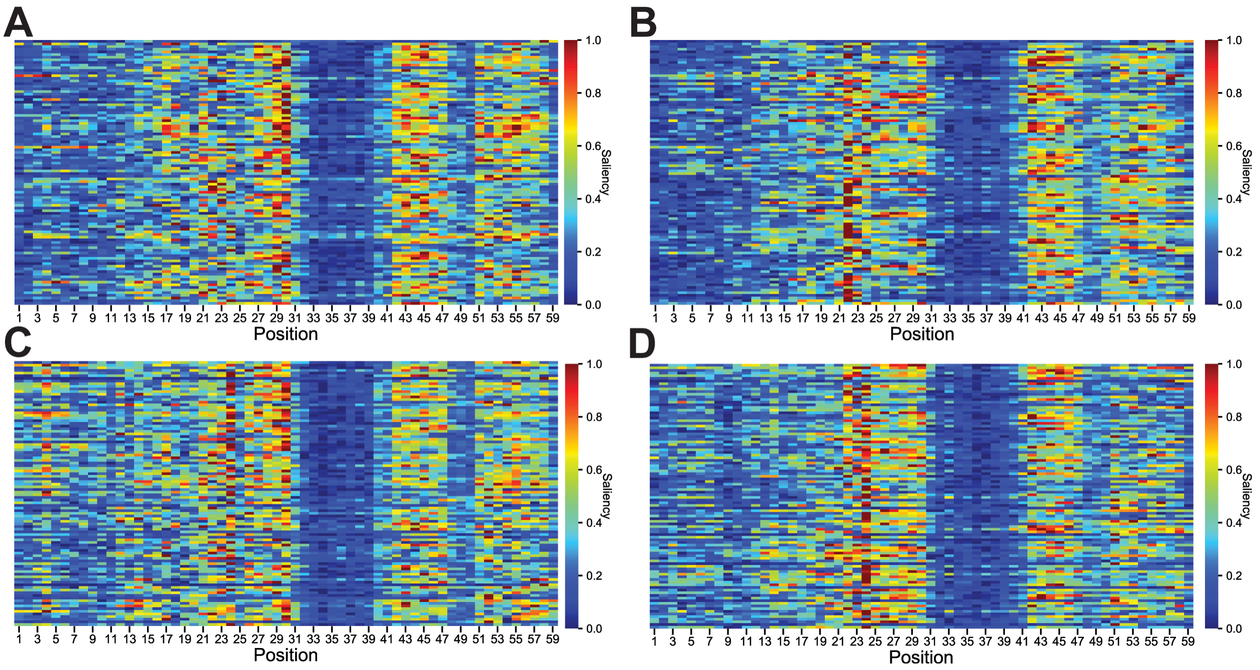


**Supplemental Figure 7: Differential attention to first twelve nucleotides in saliency maps for gradient ascent sequences pre- and post-optimization.** Given a set of 100 worst toeholds that were optimized via our gradient ascent framework, we wanted to understand how the model's focus changed when predicting ON and OFF values before and after gradient ascent. Saliency maps were generated for the pre-optimization sequences (Fig. S7A, B) and post-optimization sequences (Fig. S7C, D). For each set of sequences, both a saliency map to illustrate the parts of the sequence that maximized the ON prediction (Fig. S7A, C) as well as a saliency map to illustrate the parts of the sequence that minimized the OFF prediction (Fig. S7B, D) were generated. For most of the 100 sequences sampled, the model pays more attention to the first twelve nucleotides after optimization.

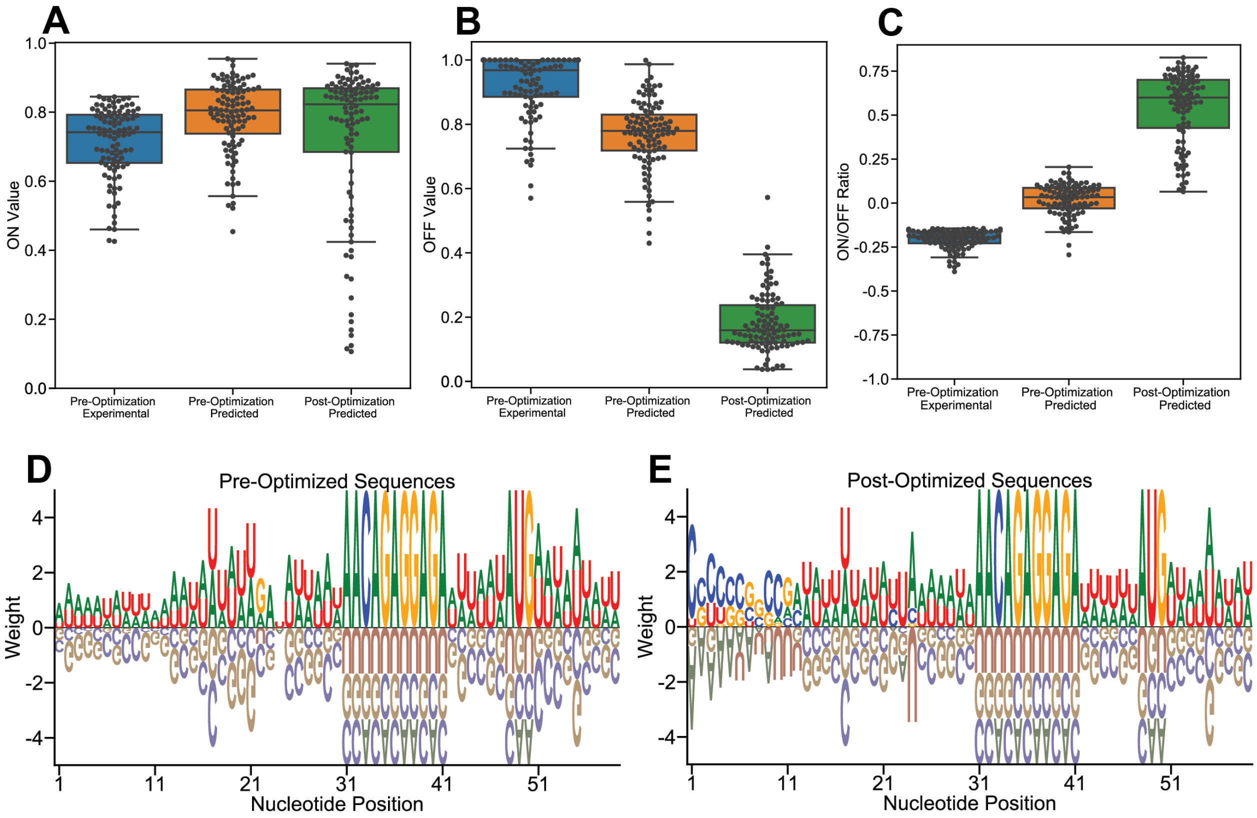


**Supplemental Figure 8: Gradient ascent on a set of toehold sequences with high correlation between experimental and predicted values results in familiar motifs.** To ensure that the results from optimizing the set of worst toehold sequences were reproducible (Fig. 3), a second set of toehold switches were optimized with our gradient ascent framework. We chose to test the gradient ascent framework on a set of 100 sequences with less than a 25% difference in experimental and predicted ON and OFF values. The experimental, predicted pre-optimization, and predicted post-optimization values for ON (Fig. S8A), OFF (Fig. S8B), and ON/OFF ratios (Fig. S8C) were plotted, with a clear increase in predicted ON/OFF ratios from pre-optimization to post-optimization. Additionally, position weight matrices for pre- and post-optimization sequences were plotted (Fig. S8D, E); though there is less consensus than in the set of worst toeholds, the run of cytosines in the first twelve positions and the over-representation of NUA at positions 22-24 as in Fig. 3E are still visible.

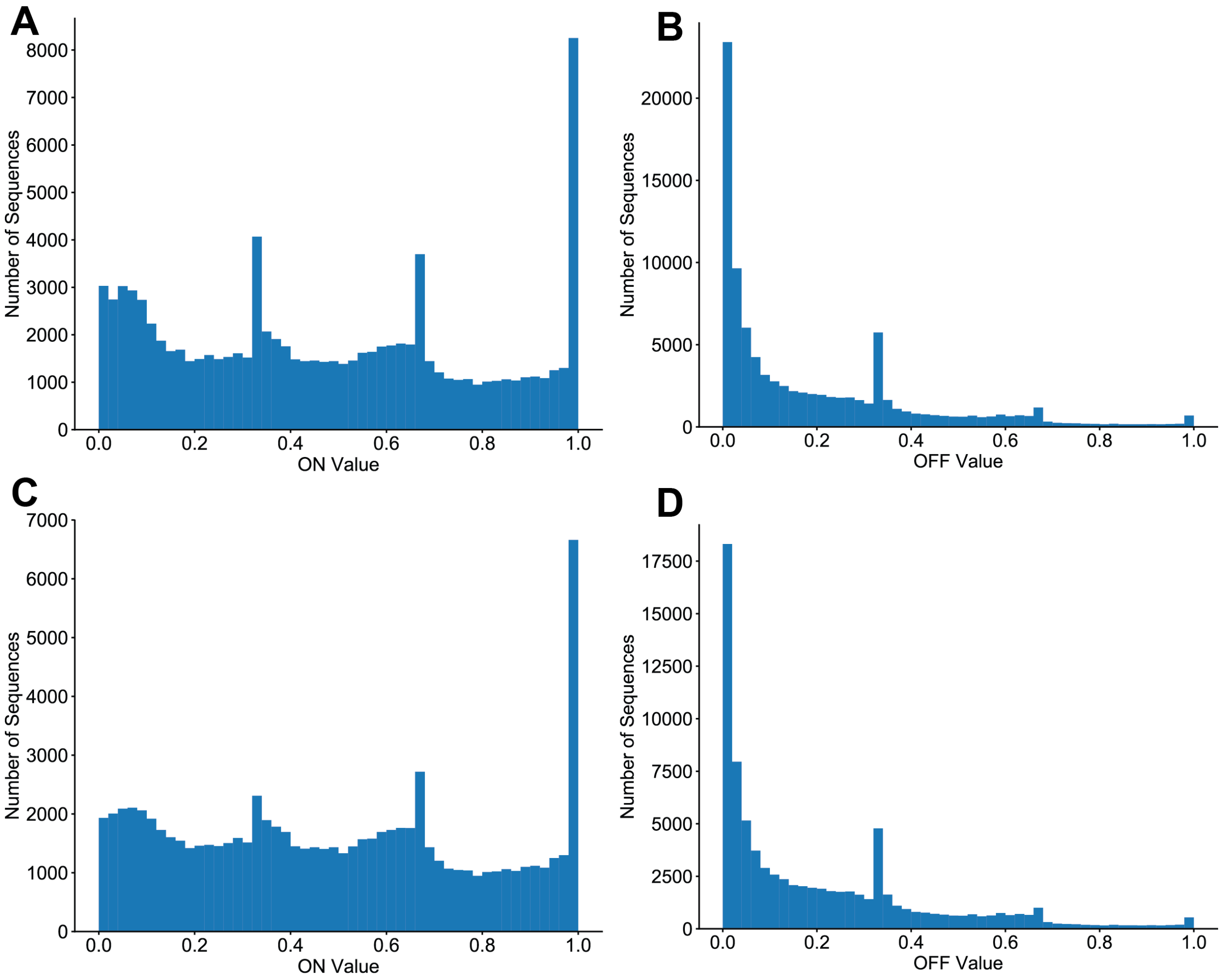


**Supplemental Figure 9: Down-sampling preserves the ON and OFF value distributions while reducing experimental artifacts.** Given the abnormally high counts observed at several ON values, the ON and OFF distributions were trimmed such that the number of sequences in each of 1000 evenly-spaced bins was reduced to the mean number of sequences over all bins. We took the union of the sequences that passed either ON or OFF filtering for a total of 81,155 sequences. Histograms of ON and OFF values were generated before (Fig. S9A, B) and after (Fig. S9C, D) down-sampling.
